# Differential Induction of Cancer Cell Death by Root, Leaf, and Flower Extract derived from *Kalanchoe pinnata*

**DOI:** 10.64898/2026.05.27.728130

**Authors:** Mitsuki Maedomari, Sakura Kawada, Nanae Harashima

## Abstract

*Kalanchoe pinnata* is a perennial plant that grows wild in tropical regions and is traditionally used as a medicinal plant. Plants of the Kalanchoe genus have been shown to possess several effects, including antibacterial and antihypertensive properties. However, effects such as the induction of apoptosis in cancer cells have not been reported for any substance other than leaf extracts of this plant and remain unexplained. Therefore, in this study, we investigated the effects of extracts from various parts of *K. pinnata* (flowers, leaves, and roots) on human colon cancer cell death. We conducted the study using three colorectal cancer cell lines (HT-29, SW620, and DLD-1) and three types of extracts derived from the flowers, leaves, and roots of this plant. Each *K. pinnata* extract significantly reduced cell viability in a dose-dependent manner in all colon cancer cells. In particular, the root extract induced cancer cell death and inhibited proliferation at lower concentrations than the other extracts. For the colon cancer cells examined, caspase-dependent apoptosis was suggested as the primary mechanism, although cell death was observed in some cells without detectable caspase activation. *K. pinnata* extracts induced both apoptosis and necrosis in colorectal cancer cells. In addition, *K. pinnata* extracts increased protein level of cleaved caspase-9, caspase-3, and PARP in SW620 and DLD-1 cells. The decrease in mitochondrial membrane potential was confirmed for all extracts, however caspase-mediated apoptosis was not observed in all cell lines, indicating the need for further investigation. Taken together, our results indicate the potential of the plant *K. pinnata* and the bioactive compounds it contains as new candidates for adjuvant therapy in colorectal cancer. In the future, it will be necessary to examine the relationship with genetic mutations in each cell line and to investigate the details of the cell death mechanism.

## 1. Introduction

Colorectal cancer (CRC) is a leading cause of cancer-related deaths worldwide; an estimated 935,000 people died from the disease in 2020, making it the second leading cause of cancer mortality after lung cancer [1]. Mortality rates for CRC are highest in Eastern Europe and East Asia; an CRC accounts for 13.7% of all cancer deaths (12.6% among men and 15.4% among women) in Japan [1]. Although chemotherapy is an effective treatment for CRC, its severe toxicity and the rapid development of drug resistance often leads to poor prognosis and reduced sensitivity to chemotherapy. Therefore, the development of potent CRC inhibitors without toxic side effects represents a promising therapeutic strategy for CRC treatment.

To prolong the survival of patients with unresectable advanced or recurrent colorectal cancer, it is essential to continue effective chemotherapy for as long as possible. Combination chemotherapy regimens such as oxaliplatin plus 5-fluorouracil (5-FU)/leucovorin (LV) (FOLFOX) and irinotecan plus 5-FU/LV (FOLFIRI) are the standard treatments for colorectal cancer, and reports indicate that long-term use of these regimens improves survival rates. However, toxicity and side effects during long-term use are significant concerns [2]. Furthermore, some cancers exhibit resistance to cytotoxic chemotherapeutic agents and molecularly targeted drugs, which can be either intrinsic or acquired. Tumors are heterogeneous populations of cancer cells with inherent genetic and functional differences, and intrinsic resistance stems from this heterogeneity. Cytotoxic chemotherapeutic agents primarily target actively proliferating cells; consequently, non-proliferating cells within the tumor—such as slow-cycle cells resulting from nutrient limitation or quiescent stem-like cells—escape treatment and cause tumor recurrence. In contrast, acquired resistance develops during anticancer drug therapy and during drug exposure to cancer cells. The mechanisms of acquired resistance are diverse and include overexpression of therapeutic targets, feedback activation of compensatory oncogenic signaling pathways, and overexpression of pumps involved in drug efflux [3]. Despite advances in chemotherapy drugs and their proven efficacy, the occurrence of side effects and the acquisition of drug resistance remain issues, creating a need for safer and more effective treatments.

The deregulation of multiple pathways, including cellular metabolism, the cell cycle, apoptosis, angiogenesis, and epigenetic modifications, which plays a significant role in the development and proliferation of cancer. Natural compounds derived from natural sources, such as plant extracts, are considered attractive candidates because they possess anticancer activity comparable to that of chemically synthesized anticancer drugs which target these pathways and may offer the potential for lower toxicity and reduced side effects [4]. Indeed, there is a growing body of research and reports demonstrating that crude extracts and constituents of natural products, such as vegetables, fruits, and medicinal plants exhibit beneficial effects against human diseases. For example, crude extracts of bitter melon have shown cancer cell proliferation-inhibiting effects accompanied by apoptosis induction in human colorectal, breast, and pancreatic cancer cells [5, 6], while crude extracts of grape skin have demonstrated similar effects in human breast and liver cancer cells [7]. Furthermore, functional components such as bromelain, a proteolytic enzyme found in pineapple [8]; cucurbitacin, found in plants of the Cucurbitaceae family [9]; and capsaicin, an alkaloid found in plants of the Capsicum genus [10], have demonstrated growth-inhibitory effects against cancer cells. Consequently, the anticancer effects of plants themselves and specific components within them are attracting significant attention. It has been reported that fucoidan, a sulfated polysaccharide derived from brown seaweed, may enable the continued administration of chemotherapy drugs in patients with unresectable, advanced, or recurrent colorectal cancer [2].

*Kalanchoe pinnata* is a perennial succulent plant belonging to the genus *Kalanchoe* in the Crassulaceae family (order Saxifragales) (Figure 1). Native to South Africa, it is distributed primarily in tropical regions and has naturalized throughout Okinawa, Japan. It has long been traditionally used in India and other countries as a medicinal plant to treat diseases such as infectious diseases, sore throat, fever, stomach pain, sprains, and wounds [11]. Regarding its relationship to cancer, there are reports that an extract from *K. pinnata* leaves induces apoptosis in HeLa cervical cancer cells via changes in the expression of the apoptosis-related factors Bax/Bcl-2 and the activation of caspase-3 and PARP-1 and suppresses HPV18 expression [12]. In this report, *K. pinnata* leaves were extracted with chloroform, and each fraction separated by column chromatography using a concentration gradient of petroleum ether and ethyl acetate demonstrated cell proliferation inhibitory effects and anti-HPV effects against cervical cancer cells [12]. In the future, if drug development is to be pursued, it is important to identify the specific plant part from which the active compounds are derived. Given the suggestion that *K. pinnata* leaf extracts possess anticancer effects [12], it is considered an essential step in drug development to investigate the anticancer effects not only of leaf extracts but also of extracts from flowers and roots.

**Figure 1.**
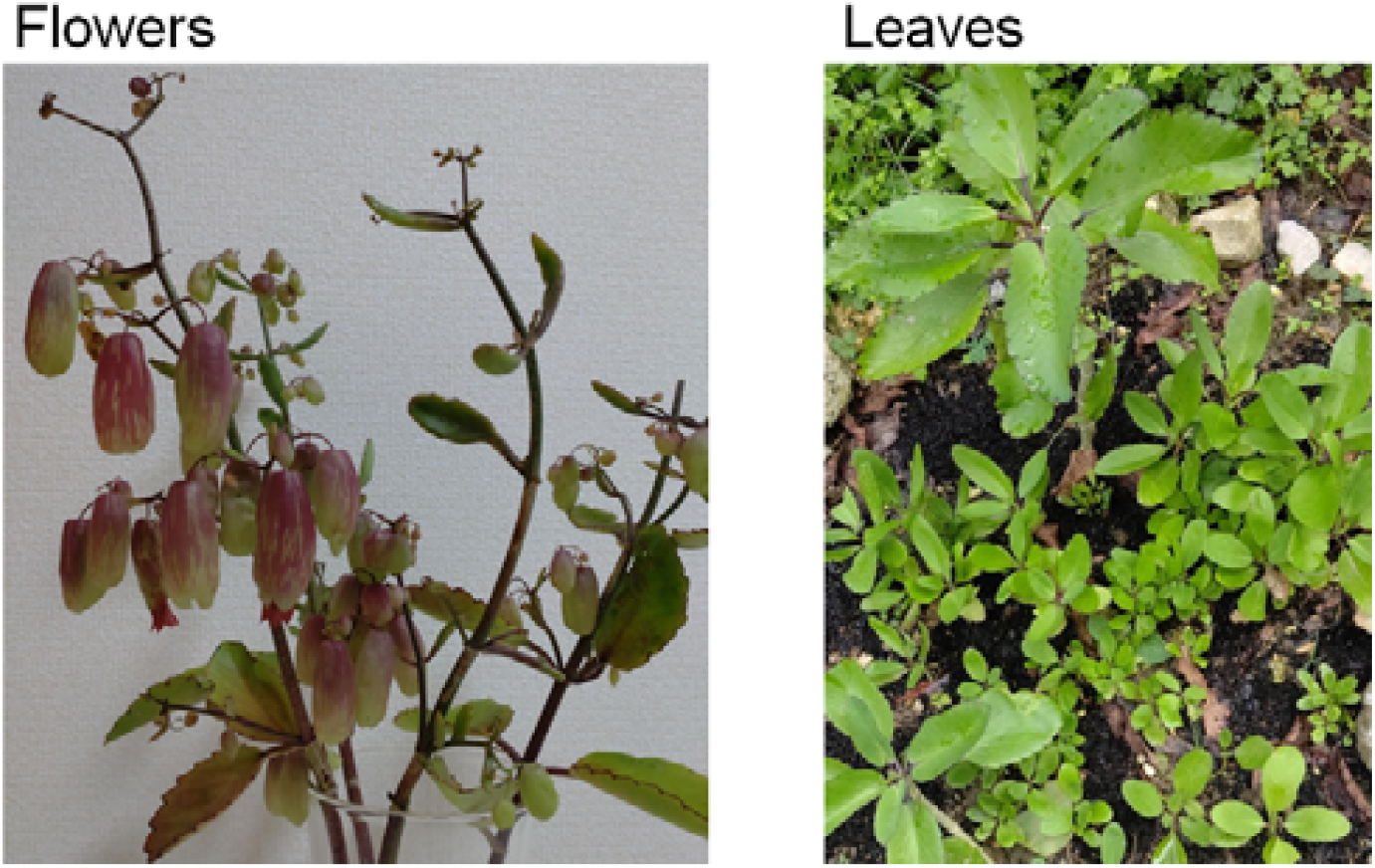
The flowers and leaves of *K. pinnata*. Photos by authors.

Therefore, in the current study, we investigated the anticancer properties of *K. pinnata* extracts from different plant parts in human colorectal cancer cells. Our data indicate that All *K. pinnata* extracts induced caspase-dependent apoptosis in cancer cells, in particular, the root extract induced cell death at a lower concentration than the others.

## 2. Materials and Methods

### 2.1. Cell culture

Human colorectal cancer cell lines (HT-29, SW620, DLD-1) were purchased from the American Type Culture Collection. Cells were maintained in DMEM containing 10% fetal bovine serum and 1% penicillin/streptomycin at 37°C with a humidified 5% CO_2_.

### 2.2. Preparation of each extract from *Kalanchoe pinnata*

*K. pinnata* (Figure 1) was obtained in Okinawa islands. The flowers, aerial parts (leaves and stems), and roots of *K. pinnata* were extracted with methanol. After removing the fat-soluble components, the constituents were adsorbed and eluted using adsorption column chromatography and finally lyophilized. The lyophilized powder was kindly provided by Dr. Kensaku Takara (Faculty of Agriculture, University of the Ryukyus) and dissolved 30 mg/mL in dimethyl sulfoxide (DMSO).

### 2.3. Cell viability

The colorectal cancer cell lines were seeded at 5 × 10^3^ cells/well in 96-well plate and cultured overnight. The working solution of *K. pinnata* extracts were added to the cells, and co-cultured for 48 hours. Next, 100 µL of culture supernatant was collected from each well, 10 µL of Cell Counting Kit-8 reagent (DOJINDO) was added, and the mixture was incubated for 3 hours. Absorbance was then measured at 450 nm using a microplate reader (SpectraMax iD5, Molecular Devises). The results were expressed as a percentage, relative to solvent control incubation. Furthermore, at lower concentrations, *K. pinnata* extracts were tested in indicated concentrations, and cultured for 48 hours before performing the same measurements. DMSO was set to 1% or less, as this has been reported to be non-toxic to cells as the solvent control in all experiments [13].

### 2.4. Cell morphological observations

The cells were seeded in 24-well plates at a density at 2 × 10^4^ cells/well. After incubation for 24 h, the cells were treated with *K. pinnata* extracts for another 48 hours. The cells morphology was observed with a phase-contrast optical microscopy.

The double fluorescent staining was performed using acridine orange (Cellstain AO, DOJINDO) and propidium iodide (Cellstain PI, DOJINDO). The cells treated with or without *K. pinnata* extracts were incubated for 30 minutes, observed under a fluorescence microscopy. After counting 100 cells per well, the average of three wells was calculated to determine the percentage of dead cells.

### 2.5. Colony Formation Assay

Colorectal cancer cells were seeded at 500 cells/well in 6-well plates and cultured overnight. *K. pinnata* extracts were added and incubated for 48 hours. The solvent control group was treated with DMSO. After 9–10 days incubation period, the cells were stained with 0.5% crystal violet, followed by a thorough wash with deionized water and air drying. The average number of colonies in the three wells containing the solvent control (DMSO) was set as 100% colony-forming ability and compared with the wells containing each *K. pinnata* extract.

### 2.7. Annexin V/PI staining

Three colorectal cancer cells were seeded at 5 × 10^5^ cells/well in a 12-well plate and incubated overnight. Flower extract was added to achieve a final concentration of 8 µg/mL for HT-29 cells and 4 µg/mL for SW620 and DLD-1 cells, leaf extract at a final concentration of 16 µg/mL, and root extract at a final concentration of 0.5 µg/mL and co-cultured for 48 hours. Then, cells were stained with Annexin V-FITC/PI (Nacalai tesque) and incubated in a dark place for 15 minutes. Subsequently, binding buffer was added to resuspend the cells, and measurements were performed using flow cytometers (MACSQUANT™ ANALYZER). Data was analyzed using FCS Express 7 (De Novo Software).

### 2.8. Mitochondrial membrane potential

The changes of mitochondrial membrane potentials induced by *K. pinnata* extracts were determined using the low-molecular-weight fluorescent dye JC-1 (MitoMP Detection Kit, DOJINDO) according to the manufacturer’s protocol. Briefly, cancer cells (5 × 10^3^ cells /well) were seeded in a 96-well plate and incubated overnight. Cells were treated with *K. pinnata* extracts for 24 hours and then incubated with JC-1 dye at 37°C for 1 hour. After washing with PBS, red fluorescence intensity was quantified using a multi-plate reader (485/535 nm) to quantify red fluorescence intensity. The red fluorescence intensity ratio for each treated group was calculated by setting the red fluorescence intensity of the solvent control group to 1.0.

### 2.9. Western Blot analysis

*K. pinnata* treated cancer cells were washed with PBS and then lysed by using M-PER Reagent (PIERCE). Proteins were separated by SDS-PAGE, transferred to a PVDF membrane, blocked with 4% skim milk/TBS-T, reacted with primary antibody (1:2000–5000) and secondary antibody (1:10,000), and then detected by chemiluminescence. The primary antibodies used were caspase-9 (Cell Signaling Technology; CST, 9508), caspase-3 (CST, 9665), PARP (CST, 9542), β-actin (abm, G043) were used as primary antibodies, and anti-rabbit IgG, HRP-linked Antibody (CST, 7074), and Anti-mouse IgG, HRP-linked Antibody (CST, 7076). Immunoreactive molecules were detected by chemiluminescence reagent Chemi-Lumi One (Nacalai tesque).

### 2.10. Statistical Analysis

All data were expressed as means ± SD for triplicate experiments. Statistical comparisons among groups were performed by Student’s *t*-test. A level of *p* < 0.05, *p* < 0.01 was considered statistically significant.

## 3. Results

### Changes in Cell Viability Following Incubation with Various *K. pinnata* Extracts

To determine whether three types of *K. pinnata* extracts influenced the cell viability of three colon cancer cells were cultured in the presence of each *K. pinnata* extract for 48 hours, and the cell viability was determined using WST-8 assay. We initially tested concentrations ranging from 25 to 200 µg/mL, but even at the lowest concentration (25µg/mL), the cell survival rate was around 30% or lower (Figure 2A). We therefore conducted tests at even lower concentrations. The cell viability in cultures treated with a final concentration of 4 µg/mL of *K. pinnata* flower extract decreased significantly to 55.8 ± 1.5 % in HT-29 cells, 26.1 ± 1.5 % in SW620 cells, and 42.4 ± 2.9 % in DLD-1 cells. In cultures with 16 µg/mL of *K. pinnata* leaf extract, cell viability decreased significantly to 50.3 ± 4.3 % in HT-29 cells, 55.0 ± 2.8 % in SW620 cells, and 60.3 ± 3.2 % in DLD-1 cells. Furthermore, in cultures with 0.5 µg/mL of *K. pinnata* root extract, viability of cancer cells was significantly reduced to 46.5 ± 12.8 %, 45.3 ± 1.5 %, and 44.0 ± 6.9 %, respectively (Figure 2B).

**Figure 2.**
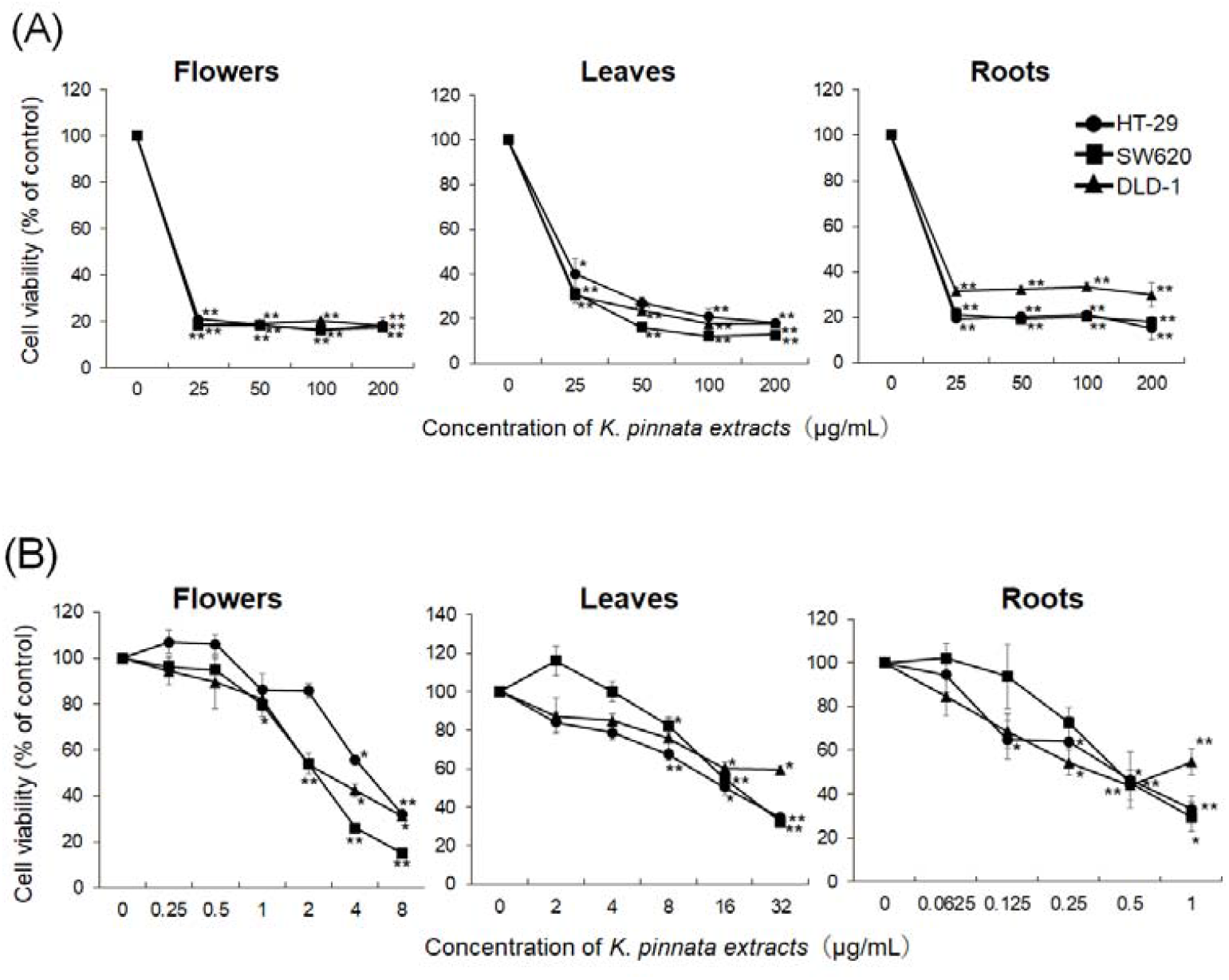
Effects of *K. pinnata* extracts in colon cancer cells. (a) Cells were treated with high concentrations of *K. pinnata* extracts for 48 hours. (b) Cells were treated with low concentrations of *K. pinnata* extracts for 48 hours. * indicates significance against the controls (\**p* < 0.05, \*\**p* < 0.01).

### Inhibition of cancer colony formation with various *K. pinnata* extracts

When three types of human colorectal cancer cells were cultured with various *K. pinnata* extracts, the colony formation ability was examined; the results showed that all *K. pinnata* extracts inhibited colony formation in all cell lines (Figure 3A). In cultures supplemented with 4 µg/mL of *K. pinnata* flower extract, the colony formation rate of HT-29 decreased to 76.0 ± 4.2 %; in cultures with 2 µg/mL, the rates of SW620 and DLD-1 decreased to 38.0 ± 13.7 % and 77.0 ±11.0 %, respectively. The rates significantly reduced to 62.3 ± 6.6 % in HT-29, 12.8 ± 10.8 % in SW620, and 46.1 ± 6.6 % in DLD-1 after treatment with 16 µg/mL of *K. pinnata* leaf extract. Furthermore, the colony formation of HT-29, SW620, and DLD-1 significantly decreased to 73.6 ± 16.2 %, 17.8 ± 10.8 %, and 16 ± 2.1 % after co-culture with 0.5 µg/mL of *K. pinnata* root extract, respectively (Figure 3B). Among these, cell proliferation in DLD-1 cells was significantly inhibited when co-cultured with root extracts, and SW620 cells showed a marked reduction in proliferation when co-cultured with any of the extracts.

**Figure 3.**
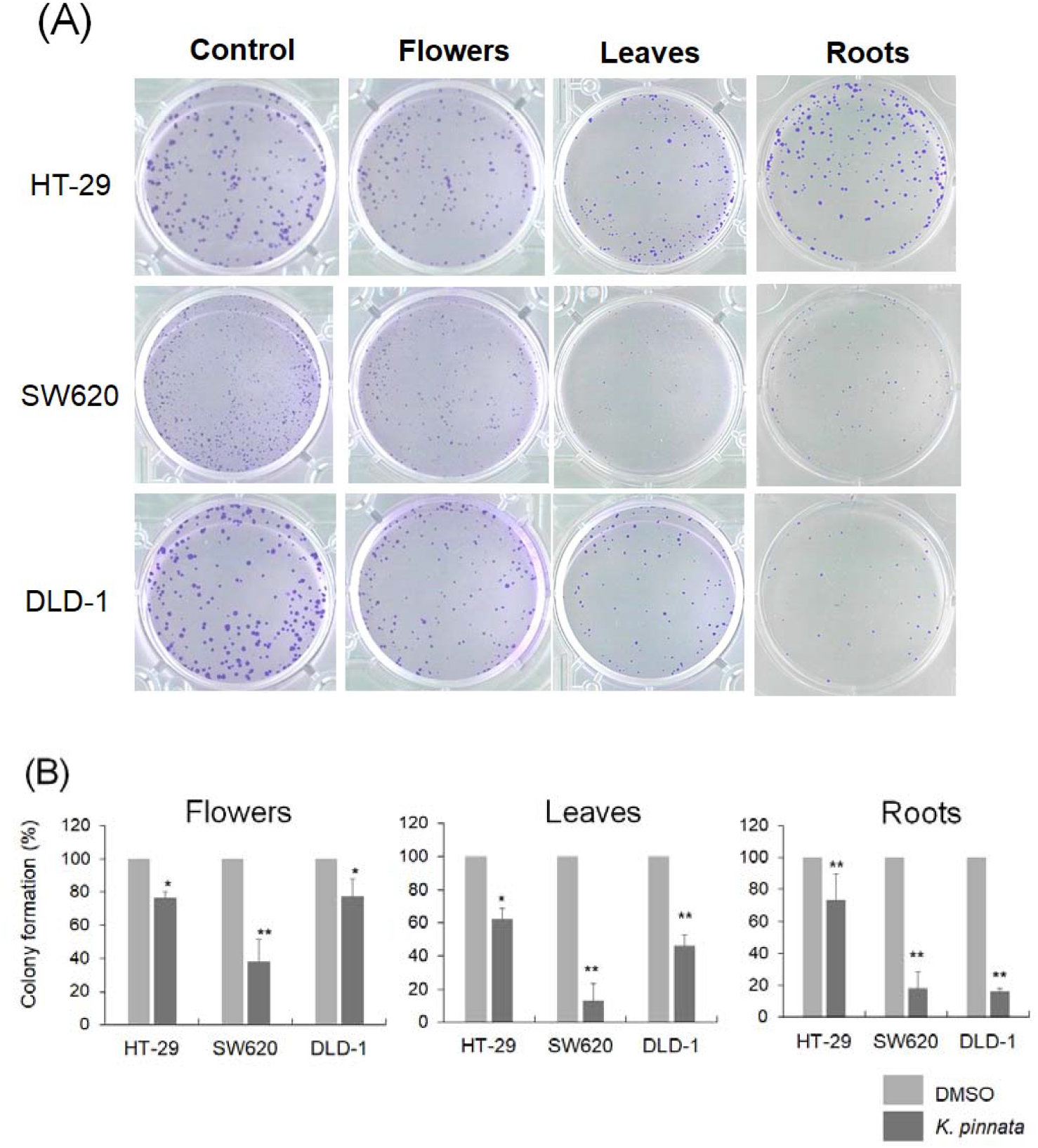
Effects of *K. pinnata* extracts on colony formation assay of three colon cancer cells. (a) Representative data of colony formation assay after treatment with K. pinnata extracts. The additional concentration was same as Figure 1 c. (b) Data are presented as the mean ± SD of the three independent wells. * indicates significance against the controls (\**p* < 0.05, \*\**p* < 0.01).

### *K. pinnata* extracts induce cell death in colorectal cancer cells

After culturing three types of colorectal cancer cells with various *K. pinnata* extracts, the cancer cells lost their adhesion and floated in the culture plates. Morphological changes of apoptosis were observed, such as widening gaps between cells and the condensation of individual cancer cells (Figure 4A). Next, cancer cells cultured under the same conditions were subjected to acridine orange (AO)/propidium iodide (PI) double staining to observe dead cells (Figure 4b). Live cells are stained green by AO, while dead cells are stained red by PI. In all three types of human colorectal cancer cells, many stained red cells were observed following culture with the various *K. pinnata* extracts. Among the cells stained red by PI, a mixture of shrunken cells, a characteristic of apoptosis and swollen, ruptured cells, a characteristic of necrosis was observed. Additionally, the red-stained cells were counted to calculate the cell death rate (Figure 4B).

**Figure 4.**
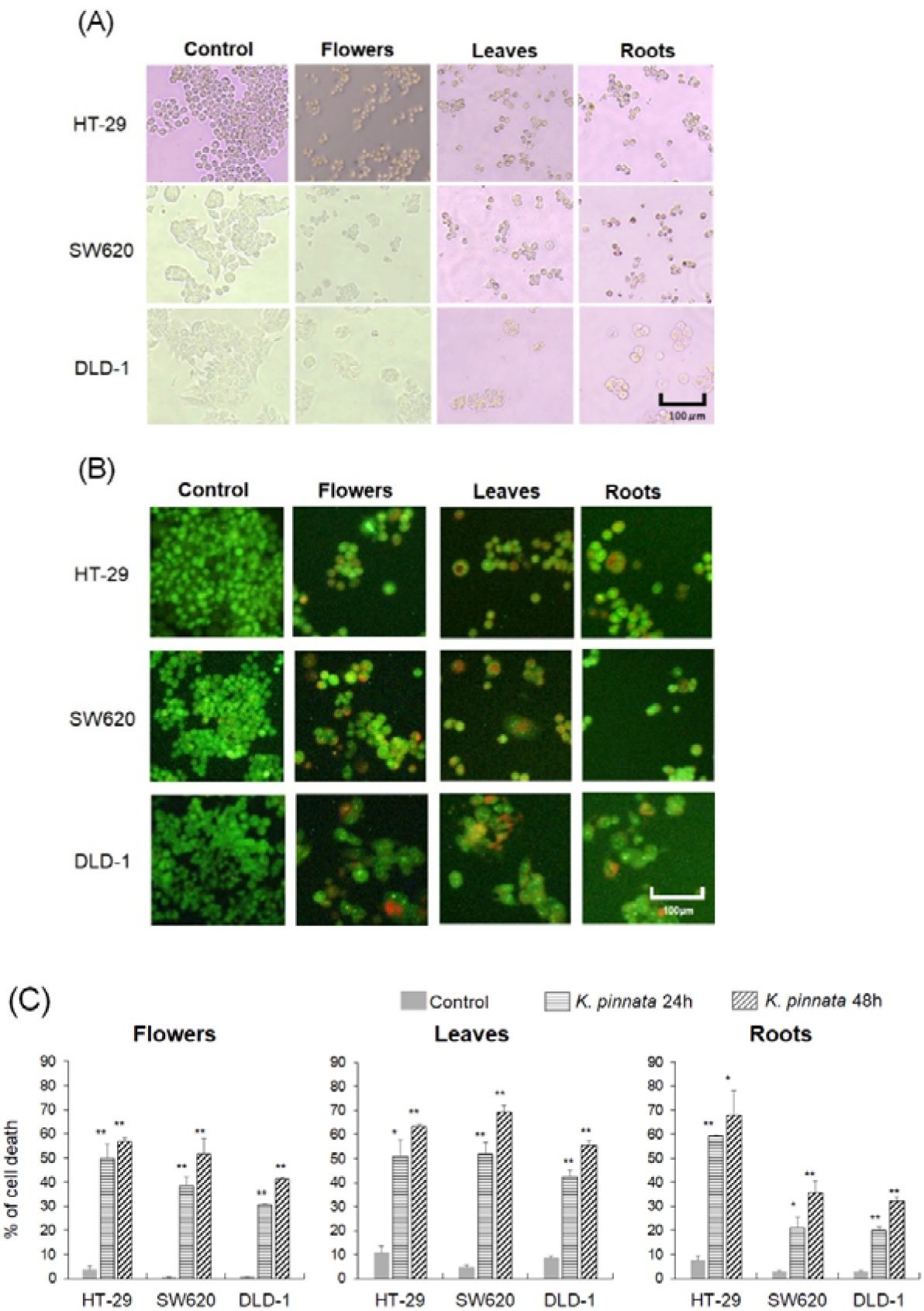
*K. pinnata* extracts induce apoptosis in colorectal cancer cells. (a) The morphological changes of the cells after treatment with *K. pinnata* extracts for 48 hours were imaged by a phase contrast microscope (magnification, 100x). (b) The cells were stained with AO/PI solutions and observed with a fluorescent microscopy (magnification, 100x). (c) After 48 hours treatment with *K. pinnata* extracts, 100 cells per well were counted and the average of three wells was calculated to determine the percentage of dead cells. Results are expressed as the mean ± SD of triplicates. * indicates significance against the controls (\**p* < 0.05, \*\**p* < 0.01).

After 24- and 48-hours co-culture with each *K. pinnata* extracts, the percent of cell death significantly increased compared to the solvent control in all tested cancer cells, and these effects were dose dependent (Figure 4C). These observations indicate that the inhibition of cell viability observed in response to *K. pinnata* extracts is also associated with the induction of cancer cell death.

### *K. pinnata* extracts induced apoptosis

After culturing three types of colorectal cancer cells with each *K. pinnata* extracts, we analyzed cell death and found no differences based on the source of the extract (Figure 5). However, the type of cell death differed among the cell lines. The Annexin V-positive cells increased in SW620 and DLD-1 cells, especially early phase of apoptotic cell population was promoted in SW620 cells after treatment with each *K. pinnata* extracts. On the other hand, the PI-positive cells, including necrotic cells was increased in treated HT-29 cells, unlike other cells.

**Figure 5.**
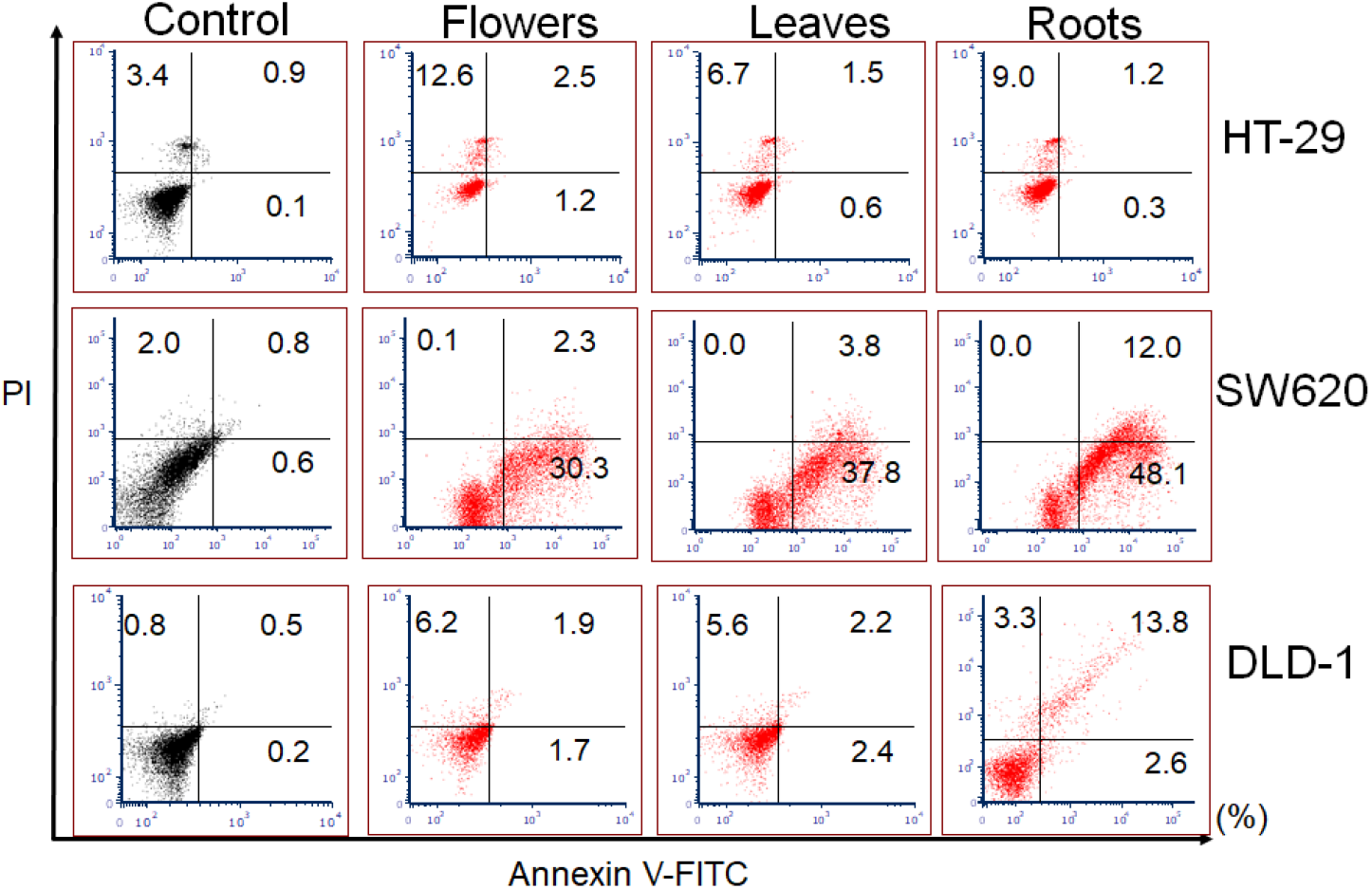
*K. pinnata* extracts induced apoptotic cell death in SW620 and DLD-1 cells. A measurement of apoptosis was determined by Annexin V-FITC and PI staining flow cytometry, which can quantitatively assess early apoptosis (lower right), late apoptosis (upper right) and necrosis (upper left).

### Activation of caspases and loss of mitochondria membrane potential

After co-culturing three types of colorectal cancer cell lines with each *K. pinnata* extracts, we examined the expression levels of caspase-9 and -3, as well as the DNA repair molecule PARP (Figure 6A). The flower extract activated caspase-9, -3 and PARP by inducing the expression of their cleaved forms in SW620 cells. The leaf extract also activated caspase-9, and -3 in SW620 and DLD-1 cells. The root extract activated caspase-9 and PARP in SW620 and DLD-1 cells. No activation of caspase or PARP was detected in HT-29 cells when co-cultured with any of the extracts. This is thought to be related to the less Annexin V staining shown in Figure 5. These results suggest that *K. pinnata* extracts induced caspase-dependent apoptosis in SW620 and DLD-1 cells, but not HT-29.

**Figure 6.**
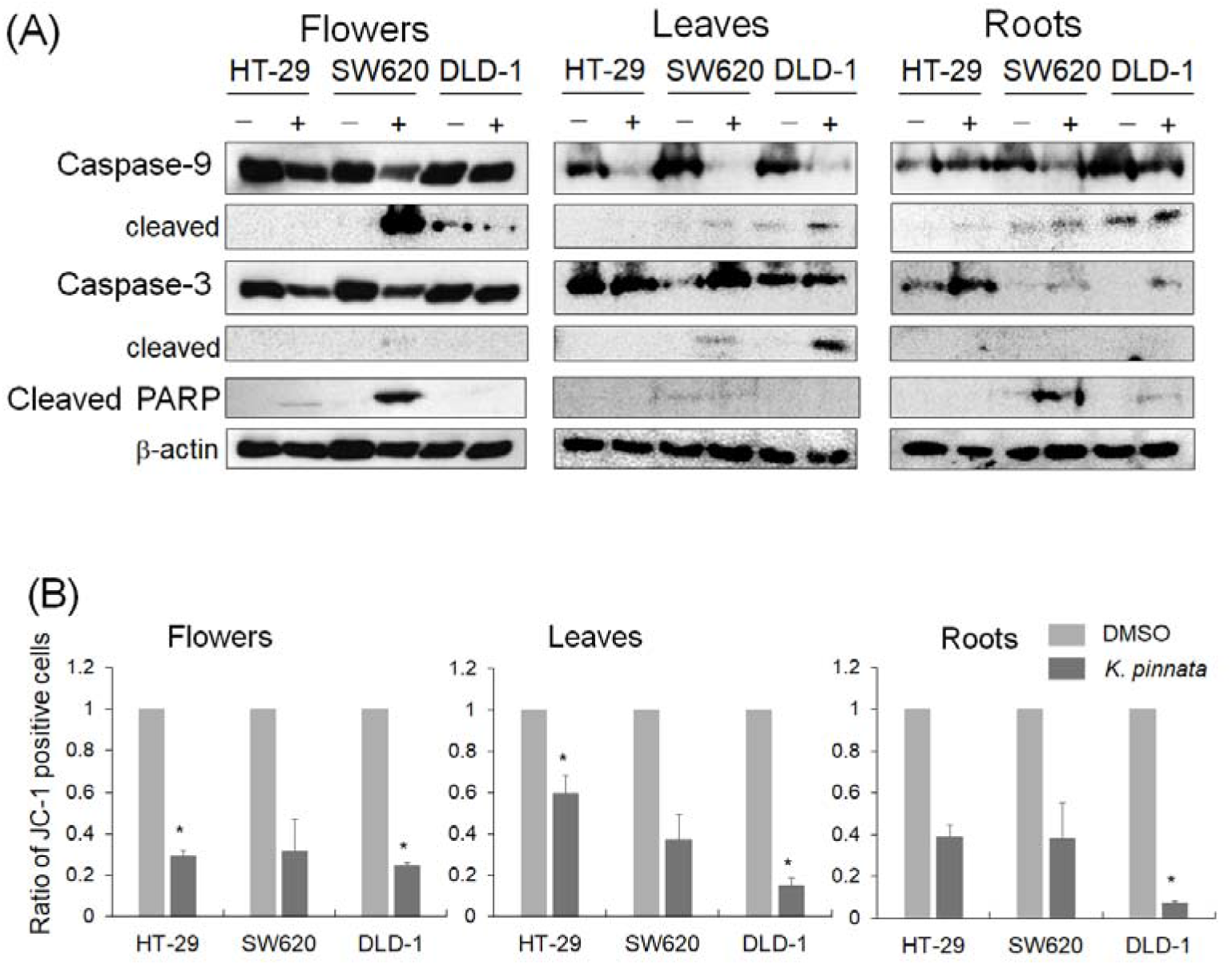
*K. pinnata* extracts stimulates apoptosis through mitochondrial pathway. (a) Colorectal cancer cells were incubated with *K. pinnata* extracts for 48 hours, then the equal amount of total cell lysates were used. The expression level of caspase-9, caspase-3, and PARP were detected by western blotting. (b)After the cells were incubated with *K. pinnata* extracts for 24 hours, the quantitative measurement of mitochondrial membrane potential was determined by JC-1 assay. Taxonomic classification (\**p* < 0.05).

We examined changes in mitochondrial membrane potential after incubating colorectal cancer cell lines with each *K. pinnata* extracts. Under normal conditions, when the membrane potential is maintained, the JC-1 dye accumulates in the mitochondria and emits red fluorescence. However, when depolarization occurs and the membrane potential decreases, JC-1 exists as monomers, and the red fluorescence intensity disappears. Since the red fluorescence intensity reduced in all three cell lines with *K. pinnata* extracts, this indicated significant decreases in mitochondrial membrane potential (Figure 6B).

## 4. Discussions

This study demonstrated that extracts from three different parts of *K. pinnata*—flowers, leaves, and roots—exhibit anticancer activity against human colorectal cancer cells *in vitro*. Each extract of *K. pinnata* inhibited the survival of colorectal cancer cells at low concentrations in culture; in particular, the root extract exhibited cancer cell growth inhibition at a very low concentration of 0.5 µg/mL (Figure 2A). Although we had expected differences in cancer cell survival rates depending on the source of the extract, no such differences were observed, all extracts inhibited cancer cell survival at low concentrations. Furthermore, each *K. pinnata* extract inhibited cell proliferation in all tested cancer cells (Figure 3). These findings suggest that each *K. pinnata* extract has a proliferation-inhibiting effect on colorectal cancer cells. The morphology of the cancer cells after co-culture with the each *K. pinnata* extracts showed reduced cell aggregation and adhesion (Figure 4A), fluorescent double staining with AO and PI revealed that PI entered the cancer cells that had undergone cell death, and numerous cells that were shrinking or swelling were observed (Figure 4B). In addition, analysis of cell death using Annexin-V/PI detected not only apoptotic cells but also necrotic cells (Figure 5). These data showed that apoptosis was induced in colon cancer cells following co-culture with *K. pinnata* extracts except HT-29 cells.

The colorectal cancer cells have previously been reported to undergo apoptosis when treated with various bioactive components or anticancer agents in combination, as evidenced by the expression of the apoptosis-initiating molecule caspase-9 and its cleaved form, the apoptosis-executing molecule caspase-3 and its cleaved form, and the DNA repair molecule PARP and its cleaved form [14-16]. To determine whether the colorectal cancer cells examined in this study were induced to undergo apoptosis, we examined the expression of cleaved caspase-9, cleaved caspase-3, and cleaved PARP. No expression of cleavage forms of caspase-9, caspase-3, and PARP were observed in HT-29 cells treated with *K. pinnata* extracts (Figure 6A). Therefore, it was considered unlikely that apoptosis was induced. The activation of cleaved caspase-9 and cleaved PARP was observed following culture with flower and root extracts, and the activation of cleaved caspase-3 was observed following culture with leaf extract in SW620 cells (Figure 6A), and then this suggested that apoptosis had occurred. Different sources derived from *K. pinnata* may be the main cause of the cell death pathway difference. In DLD-1 cells (Figure 6A), apoptosis-related enzymes were not activated by flower extract, it was considered unlikely that apoptosis was induced. On the other hand, since cleaved caspase-9 and/or caspase-3 were detected in DLD-1 cells treated with leaf or root extracts, it was suggested that apoptosis was induced and may not have affected DNA damage. Thus, it was found that even among colorectal cancer-derived cells, the cell death pathways differ depending on the cell line, and it was anticipated that this might account for differences in treatment efficacy. When cell death is induced by some stimulus and the mitochondrial membrane potential decreases, the mitochondrial permeable transition pore (MPTP) opens, allowing ions and small molecules to pass through the membrane. The resulting ionic imbalance leads to the release of cytochrome c into the cytosol, thereby inducing apoptosis [17]. Since the addition of *K. pinnata* extracts to cultures resulted in a decrease in mitochondrial membrane potential in three colorectal cancer cells (Figure 6B), it was considered that apoptosis-related molecules surrounding the mitochondria were regulated, leading to the activation of caspase-9.

Caspase-independent regulated cell death is called necroptosis, and it has recently been clarified that necroptosis signals are transmitted when caspase activity is absent, inducing exogenous apoptosis by stimulating the Fas/TNF receptor family [18]. The main signaling pathway for necroptosis involves RIP1 (receptor-interacting serine/threonine protein kinase 1) activity phosphorylating RIP3 (receptor-interacting serine/threonine protein kinase 3), which in turn phosphorylates MLKL (mixed lineage kinase domain-like protein) and simultaneously forms a trimer. Subsequently, the MLKL trimer migrates to the cell membrane, induces necrotic membrane permeabilization, and executes necroptosis [19]. The acquisition of resistance to caspase-dependent apoptosis in cancer cells poses a major obstacle in cancer treatment. Necroptosis, a form of programmed cell death, is attracting attention as a new target because it can serve as an alternative process when cancer cells exhibit resistance to apoptosis induced by chemotherapeutic agents [20, 21]. The necrotic cells were observed after treatment with *K. pinnata* extracts in HT-29 cells (Figure 5) and apoptosis-related enzymes were not activated (Figure 6) although mitochondria membrane potential was significantly decreased. Since it cannot be ruled out that apoptosis was inhibited in HT-29 cells and that necroptosis-mediated cancer cell death was induced, it will be necessary to investigate non-apoptotic cell death pathways in the future.

Anticancer drugs do not induce cell death selectively in cancer cells and exhibit toxicity to normal cells as well, making them difficult to use for long-term treatment. There is a need for new therapeutic agents that can be used for long-term treatment while suppressing side effects. As one solution to this, there are reports examining the anticancer effects on cancer cells by combining extracts derived from natural sources, such as plants, with existing anticancer drugs. An ethanol extract of lemongrass, when used in combination with FOLFOX and Taxol, induced apoptosis in human colorectal cancer cells [22], and an aqueous extract of hibiscus flowers, when used in combination with Tamoxifen, Taxol, and Cisplatin, induced apoptosis in human breast cancer cells [23], demonstrating inhibition of cancer cell proliferation. In these previous reports, apoptosis was not induced in normal cells, and cancer cell-selective cell death was observed. In many cases, plant extracts and plant-derived bioactive compounds enhance antitumor effects when used in combination with existing anticancer drugs and may block or reverse the drug resistance mechanisms acquired by cancer cells [24]. We believe that investigating whether *K. pinnata* extract is effective when used in combination with anticancer drugs will lead to expanded treatment options for cancer.

According to a previous report about *K. pinnata* [25], methanol extracts of *K. pinnata* leaves were identified as containing phenolic acids, quercetin, lycopene, β-carotene, and alkaloids. Furthermore, it was reported that phenolic acids extracted from *K. pinnata* possess high antioxidant activity and are associated with the suppression of cell survival in human acute lymphoblastic leukemia T-cell lines [26]. Since phenolic acids and quercetin have been reported to exhibit anticancer effects against cancer cells, such as apoptosis induction [27, 28], anti-angiogenesis [16, 29], and inhibition of invasion [16], it is suggested that the constituents of *K. pinnata* may influence cancer. Although there are no reports on the constituents of the flowers and roots of *K. pinnata*, they may contain components like those in the leaves or components that exhibit different anticancer effects.

In addition, various plants of the genus *Kalanchoe* contain a cardiac glycoside called bufadienolide which has been reported to possess insecticidal activity [30], antibacterial activity [31], anti-inflammatory activity [32], and cardiotonic activity [30], and is involved in numerous processes such as the induction of apoptosis and autophagy in malignant melanoma [33], as well as cell cycle arrest, anti-angiogenesis, and the inhibition of epithelial-mesenchymal transition and metastasis in various cancer cells [34]. Most of the currently reported effects of bufadienolides are derived from the parotid glands of toads or from *Kalanchoe* species other than *K. pinnata*; the anticancer effects of bufadienolides derived from *K. pinnata* have hardly been investigated. To further clarify the potential of K. pinnata as a candidate anticancer drug, it is necessary to investigate the components contained in *K. pinnata* and examine their effects on cancer cells in greater detail.

In drug discovery, it is essential that natural product-derived compounds undergo clinical evaluation by verifying the efficacy observed *in vitro* and *in vivo* animal model studies through numerous clinical trials. Furthermore, the safety of each compound when administered to cancer patients must be established [24]. In this study, we examined the anticancer effects of plant extracts, which is an early stage in the long drug discovery process. Although various studies need to be conducted in the future, the potential of plant-derived bioactive compounds is very high, and it is considered worthwhile investing them.

## 5. Conclusions

The extracts of *K. pinnata* exhibited anticancer effects against three types of colorectal cancer cells. In particular, the root extract demonstrated efficacy at significantly lower concentrations than the others. These results suggest that the *K. pinnata* extracts induce apoptosis and caspase-independent cell death, depending on the cell line used. Since the flowers, leaves, and roots of *K. pinnata* may be contained bioactive components capable of suppressing cancer characteristics such as proliferation, apoptosis, cell migration, angiogenesis, and metastasis, and regulating processes such as oxidative stress and autophagy, it will be necessary to conduct detailed compositional analyses of the three *K. pinnata*-derived extracts to identify more effective bioactive compounds in the next phase. It will also be necessary *in vivo* validation to ensure therapeutic efficacy while mitigating severe side effects and maintaining quality of life.

## Acknowledgement

The authors sincerely acknowledge Mrs. Ikuko Miyagi for her guidance and support throughout the research period. We also thank Dr. Kensaku Takara for his support preparing the plant-derived extracts. This research was supported by the University of the Ryukyus, Strategic Research Promotion Funds and a part of the project entitled “Investigation of the antitumor effects of citrus-derived flavonoids on cancer cells”, funded by Uruma Academic Research Grant Fund (2019Uruma-2400AD0301).

## Authors contribution

M.M. mainly performed the experiment, analysis of the data and drafted the manuscript. S.K. contributed reagents and analysis tools. N. H. supervised and proofread the manuscript.

